# Inhibition of glutaminase elicits senolysis in therapy-induced senescent melanoma cells

**DOI:** 10.1101/2024.09.12.612728

**Authors:** Justin Kim, Bryce Brunetti, Ayanesh Kumar, Ankit Mangla, Kord Honda, Akihiro Yoshida

## Abstract

The cyclin D1-Cyclin-Dependent Kinases 4 and 6 (CDK4/6) complex is crucial for the development of melanoma. We previously demonstrated that targeting CDK4/6 using small molecule inhibitors (CDK4/6i) suppresses Braf^V600E^ melanoma growth *in vitro* and *in vivo* through induction of cellular senescence. Clinical trials investigating CDK4/6i in melanoma have not yielded successful outcomes, underscoring the necessity to enhance the therapeutic efficacy of CDK4/6i. Accumulated research has shown that while senescence initially suppresses cell proliferation, a prolonged state of senescence eventually leads to tumor relapse by altering the tumor microenvironment, suggesting that removal of those senescent cells (in a process referred to as senolysis) is of clinical necessity to facilitate clinical response. We demonstrate that glutaminase 1 (GLS1) expression is specifically upregulated in CDK4/6i-induced senescent Braf^V600E^ melanoma cells. Upregulated GLS1 expression renders Braf^V600E^ melanoma senescent cells vulnerable to GLS1 inhibitor (GLS1i). Furthermore, we demonstrate that this senolytic approach targeting upregulated GLS1 expression is applicable even though those cells developed resistance to the Braf^V600E^ inhibitor vemurafenib, a frequently encountered substantial clinical challenge to treating patients. Thus, this novel senolytic approach may revolutionize current CDK4/6i mediated melanoma treatment if melanoma cells undergo senescence prior to developing resistance to CDK4/6i. Given that we demonstrate that a low dose of vemurafenib induced senescence, which renders Braf^V600E^ melanoma cells susceptible to GLS1i and recent accumulated research shows many cancer cells undergo senescence in response to chemotherapy, radiation, and immunotherapy, this senolytic therapy approach may prove applicable to a wide range of cancer types once senescence and GLS1 expression are induced.

**Key points:** Upregulated GLS1 expression renders senescent Braf^V600E^ melanoma cells induced by CDK4/6 inhibitor (CDK4/6i) or vemurafenib susceptible to GLS1 inhibitor (GLS1i) even with Braf^V600E^ inhibitor resistance. This senolytic therapy combining CDK4/6i and GLS1i provides insights into potential novel therapeutic strategies for metastatic melanoma and may be applicable to various types of cancers providing alternative therapy options.

## Introduction

Senescence is a state of stable and irreversible cell cycle arrest that occurs as a natural part of aging, as well as in response to various cellular stressors ^1–4^. It is an essential cellular process that plays a crucial role in maintaining tissue homeostasis and preventing the proliferation of damaged or dysfunctional cells. In recent years, senescence has emerged as a mechanistic double-edged sword for cancer. While the loss of proliferative capacity of senescent cells appears to prevent the accumulation of aberrant cells, thus acting as a barrier to tumorigenesis, senescent cells secrete a variety of inflammatory cytokines and form chronic microinflammation in the surrounding tumor microenvironment, promoting carcinogenesis, resistance to tumor immunity, and metastasis ^5–8^. Furthermore, senescent cancer cells can stimulate the growth of neighboring non-cancerous cells such as stromal fibroblasts, potentially contributing to therapy resistance and tumor relapse ^9–11^. Senolytic therapy is an emerging therapeutic approach that specifically targets and eliminates senescent cells, which prevents cancer recurrence ^12–15^ as well as extending a healthy lifespan in murine models ^16, 17^. Incorporating senolytic therapy into existing cancer treatment regimens has the potential to significantly improve patient outcomes and transform cancer management ^18, 19^.

Advanced metastatic melanoma, primarily driven by Braf (∼50%) (primarily Braf^V600E^) and Nras (∼20%) mutations, portends a dismal prognosis, with a 5-year overall survival rate of less than 50% ^20, 21^. Despite advancements in immuno- and targeted therapies ^22^, the prognosis remains poor for the subset of melanoma patients who do not respond to immune checkpoint inhibitors, representing an unmet clinical need. Dysregulated cell cycle progression, driven by Cyclin-Dependent Kinases 4 and 6 (CDK4/6) and their catalytic subunits D type cyclins, is a hallmark of cancers ^23–25^. We demonstrated that CDK4/6 inhibitor (CDK4/6i) suppressed Braf^V600E^ melanoma proliferation through senescence induction ^26–28^. However, FDA-approved CDK4/6is for ER-positive/HER2-negative breast cancer, have had modest success in melanoma clinical trials^29^.

Previous research demonstrated that cancer cells exhibit increased glutamine metabolism, which fuels the tricarboxylic acid (TCA) cycle, nucleotide and fatty acid biosynthesis, and redox balance in cancer cells ^30, 31^. This glutamine addiction is frequently observed in several types of human cancers and is driven by multiple oncogenes such as cyclin D1 or Myc ^32–34^. Therefore, targeting glutamine metabolism is a novel strategy for cancer treatment. The antitumor activity of CB-839, a glutaminase inhibitor, has been the subject of several clinical trials (clinicaltrials.gov- NCT02071862, NCT02771626), but these trials have been unsuccessful, suggesting that additional strategies that enhance the treatment efficacy are of clinical necessity. Recent studies demonstrated that CDK4/6i reprograms mitochondria metabolism, which elicits unique vulnerabilities ^35, 36^. Glutaminase 1 (GLS1) is an essential gene for the survival of human senescent cells and inhibition of GLS1 eliminated senescent cells resulting in improved age-associated organ dysfunction ^37^. Importantly, senolysis following cancer therapies has been postulated as an attractive approach to eliminating cancer cells as well as extending a healthy lifespan. However, it is unclear whether or not GLS1 is essential for the survival of senescent cancer cells induced by therapies such as CDK4/6i.

In this study, we demonstrate that prolonged treatment of Braf^V600E^ melanoma cells with the CDK4/6i induces the expression of KGA, a major isoform of GLS1, in Braf^V600E^ melanoma cells, but not in Nras^Q61^ melanoma cells. CDK4/6i-induced senescent cells become sensitive to GLS1 inhibition and undergo cell death *in vitro* and *in vivo*. We further demonstrate that the senolytic therapy is applicable for Braf^V600E^ inhibitor vemurafenib-resistant melanoma cells, suggesting this strategy can be used after patients develop resistance to Braf^V600E^ targeted therapy. Together, these results indicate that this sequential combinatorial therapy approach using CDK4/6i and GLS1i may improve the current landscape of treating patients with Braf^V600E^ melanoma.

## Materials and Methods

### Tissue culture

Authenticated human melanoma cell lines (WM3918, WM3912, A375, 1205Lu, and WM3451) were obtained from Dr. Meenhard Herlyn at the Wistar Institute (Philadelphia, PA). Human embryonic kidney (HEK) 293T cells, SK-MEL-28. SK-MEL-2, and Primary Epidermal Melanocytes; Normal, Human, Adult (HEMa) were obtained from ATCC. WM3918, WM3912, A375, 1205Lu and SK-MEL-28 were cultured in Dulbecco’s modified Eagle’s medium (DMEM) supplemented with 2% fetal bovine serum (FBS), 100 U/mL penicillin, and 100 μg/mL streptomycin. HEK 293T cells were cultured in DMEM with 10% FBS, penicillin (100 U/mL), and streptomycin (100 μg/mL). SK-MEL-2 and WM3451 cells were cultured in MCDB153 medium supplemented with 20% Leibovitz L-15, 2% FBS, 1.68 mM CaCl2, 100 U/mL penicillin, and 100 μg/mL streptomycin. HEMa (PCS-200-013) cells were cultured in Dermal Cell Basal Medium (PCS-200-030) supplemented with Adult Melanocyte Growth Kit components (PCS-200- 042), Penicillin-Streptomycin-Amphotericin B Solution (PCS-999-002) and Phenol red (PCS-999- 001).

### Plasmids and viral production

HEK293T cells were co-transfected with either GLS shRNA-encoding or KGA overexpression plasmids along with packaging plasmids (pMD2.G and psPAX2 or Qψ) using Lipofectamine 2000 (Invitrogen). Viral supernatants were collected and filtered 72 hours post-transfection and used to infect melanoma cells in the presence of 1 μg/mL polybrene (Millipore). Selection of stably transduced cells was achieved by applying 0.5 to 1 μg/mL puromycin or 1 mg/mL G418 to the growth medium, with antibiotic treatments administered every 2 to 3 days over a total of 8 days. For constructs with inducible expression, transduced cells were treated with 1 μg/mL doxycycline (Thermo Fisher Scientific cat# J60422-06) for 5 days. Transduction efficacy was confirmed by Western blotting or qRT-PCR analysis. TRIPZ inducible lentiviral human GLS shRNA clones (V2THS_233550 with the sequence 5’-TTAACAACTTCAACATGAC-3’ and V3THS_405623 with the sequence 5’-AAAATATACATCACTACCA-3’) were obtained from Dharmacon Reagents. The pLKO.1_puro control vector was provided by Dr. Bob Weinberg (Addgene plasmid #8453). pQC hKGA.wt V5-His IRES G418 (Addgene plasmid #110390), tet-pLKO.puro_shGAC (Addgene plasmid #110423), and pLKO.1puro_shGLS (Addgene plasmid #110335) were gifts from Dr. Sandra Martha Gomes Dias, with shRNA sequences as follows: 5’- CCTCTGTTCTGTCAGAGTT-3’ for shGAC and 5’-CAACTGGCCAAATTCAGTC-3’ for shGLS.

### Reverse transcription-quantitative polymerase chain reaction (RT-qPCR)

Total RNA was isolated utilizing the RNeasy Mini Kit (Qiagen) followed by reverse transcription with the iScript cDNA Synthesis Kit (Bio-Rad) as per the protocols provided by the supplier. The RT-qPCR assays were conducted using the SsoAdvanced Universal SYBR Green Supermix (Bio-Rad), with gene expression levels of KGA, GAC, and GLS2 normalized to the reference gene glyceraldehyde-3-phosphate dehydrogenase (GADPH) or beta-actin (ACTB). The sequences of primers are as follows: 5’-ACT GGA GAT GTG TCT GCA CTT CGA AG-3’ and 5’-CCA AAG TGC AGT GCT TCA TCC ATG GGA GTG-3’ for KGA, 5’-TTG GAC TAT GAA AGT CTC CAA CAA GAA C-3’ and 5’-CCA TTC TAT ATA CTA CAG TTG TAG AGA TGT CCT C-3’ for GAC, 5’-TGA GGC ACT GTG CTC GGA AGT T-3’ and 5’-TCG AAG AGC TGA GAC ATC GCC A-3’ for GLS2, 5’-CAG CCT CAA GAT CAT CAG CA-3’ and 5’-GTC TTC TGG GTG GCA GTG AT-3’ for GAPDH, 5’-CTC TTC CAG CCT TCC TTC CT-3’ and 5’-AGC ACT GTG TTG GCG TAC AG-3’ for ACTB.

### Western blot

Cells were lysed using cOmplete™ Lysis-M, an EDTA-free reagent (Roche, cat# 04719964001), to which a cOmplete™ Mini, EDTA-free protease inhibitor cocktail (Roche, cat# 04693159001) was added. The lysates were centrifuged at 15,000 rpm for 30 minutes at 4°C, after which the supernatants were collected. Protein concentrations were quantified using the Bradford assay, with the protein assay dye reagent concentrate (Bio-Rad, cat# 5000006). The lysates were then combined with sample buffer and heated at 98°C for 7 minutes. Equal amounts of protein were separated by SDS-polyacrylamide gel electrophoresis (SDS-PAGE) and then electro-transferred onto PVDF membranes. The membranes were blocked in 5% BSA for 2 hours at room temperature and incubated overnight at 4°C with the following primary antibodies: anti-glutaminase-1/GLS1 (E9H6H), anti-beta-actin (8H10D10), anti-phospho-histone H2A.X (Ser139) (20E3), anti- phospho-S6 ribosomal protein (Ser235/236) (2F9), anti-S6 ribosomal protein (5G10), anti- phospho-p44/42 MAPK (Erk1/2) (Thr202/Tyr204) (D13.14.4E), and anti-p44/42 MAPK (Erk1/2) (137F5) obtained from Cell Signaling Technology. All primary antibodies were diluted in TBST with 5% BSA to the manufacturer’s recommended concentrations. Following incubation with the primary antibodies, the membranes were washed and incubated with horseradish peroxidase (HRP)-conjugated secondary antibodies suitable for the primary antibodies used, either anti-mouse or anti-rabbit HRP-conjugated secondary antibodies (Cell Signaling Technology), for one hour at room temperature. The secondary antibodies were prepared in non-fat milk according to the manufacturer’s instructions. After additional washing steps, the bound antibody complexes were visualized using enhanced chemiluminescence (ECL) detection reagents (Bio-Rad, cat# 1705061). The immunoreactive bands were captured using autoradiography. The obtained images were scanned and bands were quantified using ImageJ software.

### Annexin V staining

Cultured melanoma cells with or without treatments were harvested and washed twice with PBS. The cell pellet was resuspended in 1x binding buffer at a concentration of 1×10^6^ cells/ml. Subsequently, 100 μl of the suspension (1×10^5^ cells) was incubated with 5 μl of APC Annexin V (BD Biosciences) and incubated for 15 minutes at room temperature. Cells were analyzed by flow cytometry.

### Immunohistochemistry (IHC)

Mouse xenograft tissue samples were fixed in 10% formalin, followed by a series of ethanol washes for deparaffinization and rehydration. Tissue sections on slides were subjected to blocking with 10% goat serum before incubation with primary antibodies. The IHC protocol included the application of biotinylated rabbit secondary antibodies, visualization with an ABC substrate kit, and subsequent development with a DAB chromogen, all from Vector Laboratories. Counterstaining was performed using hematoxylin (Thermo Fisher Scientific), with slides mounted using Permount (Thermo Fisher Scientific). The antibodies used in the study is as follows: anti-glutaminase-1/GLS1 (E9H6H) antibody and anti-cleaved caspase-3 (Asp175) antibody obtained from Cell Signaling Technology and rabbit secondary antibody conjugated to biotin obtained from Sigma Aldrich. The staining intensity was assessed on a scale from 0 (no staining) to 3 (strong staining). The percentage of positive cells was estimated on a scale from 0 (no population) to 3 (100% positive). Each specimen was scored by multiplying the intensity by the percentage of positive cells, yielding a raw score. The H-score was calculated for each section using the formula: H = Σ (I × P), where I represents the intensity scores, and P represents the percentage of positive cells.

### Senescence-Associated β-Galactosidase (SA-β-gal) Staining

SA-β-gal activity was assessed in both in vitro cultured melanoma cells and in vivo melanoma tumor sections embedded in Tissue-Tek O.C.T. compound (Sakura Finetek). Following treatment with drugs, both *in vitro* and *in vivo* samples were fixed and stained using the X-gal staining solution (Sigma) following the manufacturer’s guidelines.

### Xenograft experiments

A total of 1 × 10^6^ melanoma cells were subcutaneously injected into 6-week-old SCID mice with Matrigel (BD Biosciences). Mice were treated with vehicle, palbociclib (90 mg/kg), or CB-839 (200 mg/kg) by oral gavage once a day for 17 days after tumors had developed 5 mm in diameter following inoculation. For the combination group, mice were treated with palbociclib for 8 days and then treated with palbociclib and CB-839 for an additional 9 days. Tumor volumes were measured by caliper every 2 days and calculated by the following formula: V = length × width × width / 2. Mice were euthanized, and the tumor size was measured at day 17. Experimental animal maintenance was conducted in compliance with the institutional guidelines of Case Western Reserve University.

### Quantification of mitotic index and necrotic cells

The hematoxylin and eosin (H&E)-stained slides of all tumors were scanned using Aperio AT2 imager (Leica) and utilized for analysis of mitotic index and necrotic proportion. The mitotic cells were enumerated within a representative 1 mm^2^ area, and the proportion of necrotic cells was quantified in each scanned H&E image using QuPath software.

### Statistical analysis

Parametric data were expressed as mean ± standard deviation (SD), and non-parametric data were expressed as median and interquartile range (IQR). To compare the two independent groups, Student’s t-test was used for parametric data, and the Mann-Whitney U test was employed for non-parametric data. Experiments were performed at least three times independently. Statistical significance was set at a p-value of less than 0.01. All analyses were performed using GraphPad Prism.

## Results

### GLS1 specifically KGA is upregulated in senescent Nras WT melanoma cells

Glutaminase (GLS1) is an enzyme known to convert glutamine to glutamate in the Tricarboxylic acid (TCA) cycle. GLS1 consists of two splice variants, Kidney-type glutaminase (KGA) and glutaminase C (GAC) ^38^. KGA and GAC are identical except for amino acids in the c-terminus. GLS2, referred to as liver-type glutamine, is derived from a gene distinct from GLS1. To determine the expression of KGA, GAC and GLS2 in senescent cells induced by prolonged treatment of CDK4/6i, we treated a variety of melanoma cell lines harboring different oncogenic drivers (Braf^WT^Nras^WT^, Braf^V600E^Nras^WT^ (Braf^V600E^) and Braf^WT^Nras^Q61R/K^ (Nras^Q61^)) with CDK4/6i palbociclib for 8 days, compared to normal human melanocyte (ATCC-HEMa) as a control. Importantly, we previously demonstrated that while CDK4/6i treatment for 1 day induces reversible cell cycle arrest (quiescence), prolonged CDK4/6i treatment for 8 days induces senescence in Braf^V600E^ melanoma cell lines ^26, 27^. We demonstrated that palbociclib induced senescence specifically in Braf^WT^Nras^WT^ and Braf^V600E^ melanoma senescent cells but not in Nras^Q61R/L^ cell lines nor primary human melanocytes as assessed by SA-βgal assay, an assay to determine senescence induction (Fig 1A and Sup Fig 1). Senescence was not induced in Nras^Q61^ melanoma cell lines by palbociclib as Nras^Q61^ mutant activates both MAPK signaling and PI3K- mTOR pathway and we previously demonstrated that mTORC1 signaling activation is the resistance mechanism to CDK4/6i ^26^. Therefore, cells harboring Nras^Q61^ are refractory to CDK4/6i- induced senescence. Quantitative PCR (QPCR) gene expression analysis revealed that KGA expression is induced only in 8 days palbociclib treatment-induced senescent cells but not 1-day palbociclib treatment-induced cell cycle arrested cells (Fig 1B). The expression of GAC and GLS2 are marginally changed, but these changes are not as robust as KGA upregulation (Fig 1C-D).

**Figure 1.**
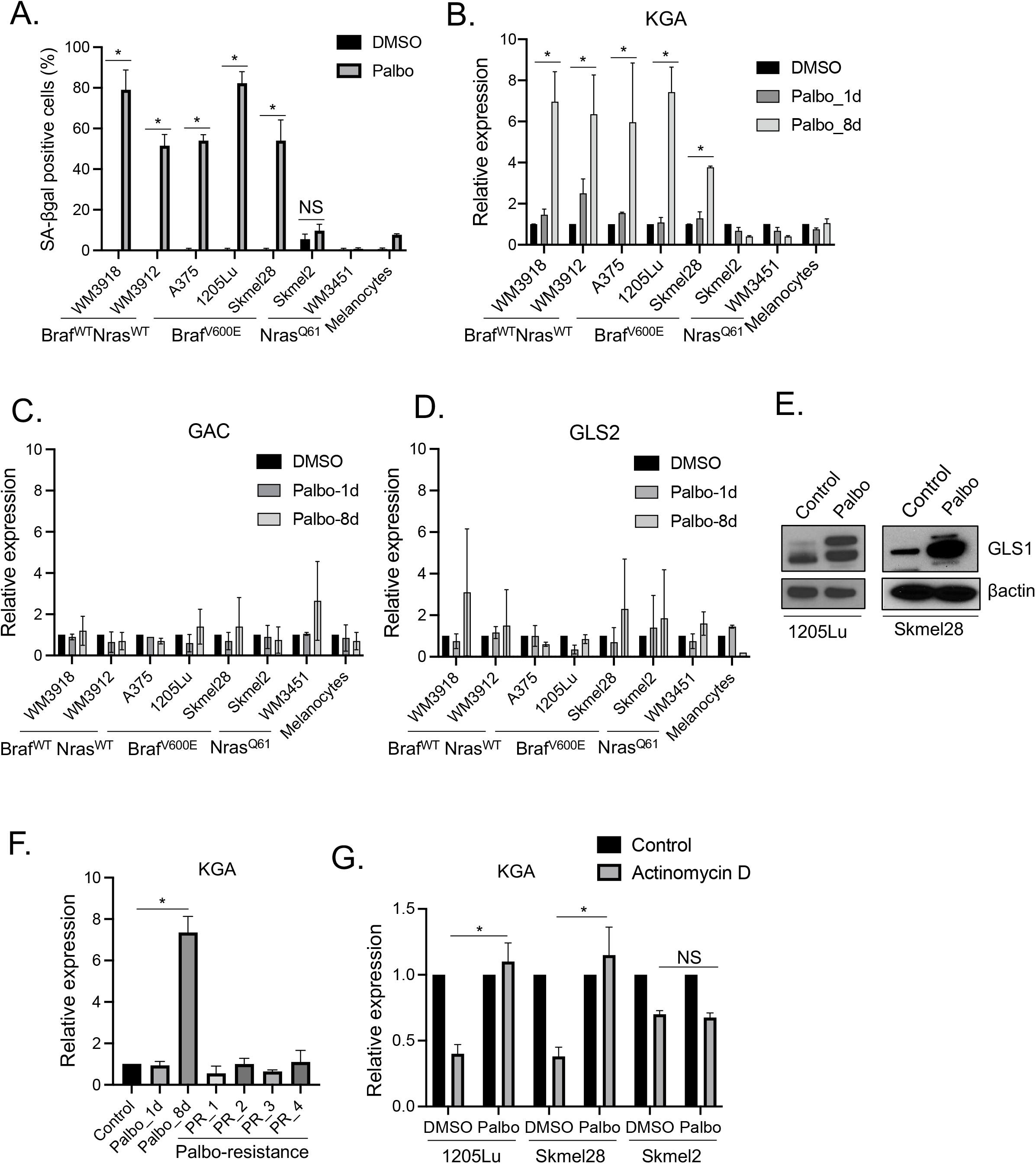
KGA expression is increased in palbociclib-induced senescent cells. (A-D) WM3918 (Braf^WT^Nras^WT^), WM3912 (Braf^WT^Nras^WT^), A375 (Braf^V600E^Nras^WT^), 1205Lu (Braf^V600E^Nras^WT^), Skmel28 (Braf^V600E^Nras^WT^), Skmel2 (Braf^WT^Nras^Q61R^), WM3451 (Braf^WT^Nras^Q61K^), and normal human melanocytes (HEMa) were treated with DMSO or palbociclib (1μM) for 1 day or 8 days. Harvested cells were subjected to SA-βgal assay (8 days treatment only) (A) and RT-qPCR analysis for KGA (B), GAC (C) and GLS2 (D). RT-qPCR data (B-D) were normalized by GAPDH and represent mean ± SD, * *p* < 0.05 (two-tailed Student’s t- test; n = 3). (E) 1205Lu and Skmel28 cells treated with DMSO (Control) or palbociclib (1μM) for 8 days were subjected to western blot for GLS1 and βactin. (F) 1205Lu cells treated with DMSO (Control) or palbociclib (1μM) for 1 day or 8 days and 4 clones of 1205Lu palbociclib resistant cells (PR_1, PR_2, PR_3 and PR_4) were harvested and subjected to RT-qPCR analysis for KGA expression. Data were normalized by GAPDH and represent mean ± SD, * *p* < 0.05 (two-tailed Student’s t-test; n = 3). (G) 1205Lu, Skmel28 and Skmel2 cells were treated with DMSO or palbociclib for 8 days, and then treated with actinomycin D (0.5 μg/ml) for 2 hours. Harvested cells were subjected to RT-qPCR analysis for KGA expression. Data were normalized by GAPDH and represent mean ± SD, *p* < 0.05 (two-tailed Student’s t-test; n = 3). NS indicates no significance (n = 3).

Consistent with a previous study ^39^, primary human melanocytes did not undergo senescence following CDK4/6i treatment because normal human melanocytes are not dependent on CDK4/6 to proliferate compared with melanoma cell lines (Fig 1A-B). Upregulation of KGA after palbociclib treatment for 8 days in Braf^V600E^ melanoma cell lines was verified by western blot analysis (Fig 1E). We previously established four independent clones of palbociclib and ribociclib-resistant (PR) cells from 1205Lu Braf^V600E^ melanoma cells ^26^. We used those PR cells to assess the expression of KGA and demonstrated that KGA is not upregulated (Fig 1F) compared to control cells while KGA is upregulated only in palbociclib-induced senescent cells, underscoring the specificity of KGA upregulation in CDK4/6i-induced Braf^V600E^ senescent cells. Mechanistically, it has shown that the GLS1 transcript contains an AU-rich element (ARE) within the 3′ untranslated region (3′UTR) ^40^, which increases mRNA stability through RNA binding protein HuC binding under acidic conditions ^41, 42^. The GLS1 transcript was stable in senescent cells but unstable in non-senescent cells and HuC depletion in senescent cells reduced survival of senescent cells ^37^. Therefore, we assessed the mRNA stability of KGA in palbociclib-induced senescent cells using the transcription inhibitor actinomycin D. We treated Braf^V600E^ 1205Lu and skmel28 cells and Nras^Q61R^ skmel2 with or without palbociclib for 8 days followed by actinomycin D treatment for 2 hours to assess the expression of KGA by QPCR. While KGA mRNA is reduced by approximately 50% in control cells after 2 hours of actinomycin D treatment, KGA mRNA is stabilized in palbociclib-induced Braf^V600E^ melanoma senescent cells (Fig 1G). In contrast, KGA expression is unchanged by actinomycin D before and after treatment of CDK4/6i in Nras^Q61R^ skmel2 cells (Fig 1F), demonstrating the specificity of KGA upregulation in Braf^V600E^ melanoma cells.

### GLS1 inhibitor induces senolysis in Nras WT senescent melanoma cells

GLS1 upregulation is essential for the survival of senescent cells because glutaminolysis produces ammonia, which neutralizes the acidic conditions caused by the leaking of intralysosomal H+ from damaged lysosomes in these senescent cells. However, when GLS1 is inhibited, senescent cells cannot produce ammonia to neutralize the acidic pH, leading to cellular acidosis ^37^. We next assessed whether the specific upregulation of KGA expression in palbociclib-induced senescent cells renders Braf^V600E^ melanoma cells vulnerable to GLS1 inhibitor (GLS1i). We treated a variety of melanoma cell lines (Braf^WT^Nras^WT^, Braf^V600E^, and Nras^Q61R/L^) with palbociclib for 1 day or 8 days, followed by a specific GLS1i (CB-839) for an additional 3 days to assess cell death by Annexin V staining, a well-established marker for apoptosis (Fig 2A). We demonstrated that inhibition of GLS1 by CB-839 eliminated 30-50% of palbociclib-induced senescent Braf^WT^Nras^WT^ and Braf^V600E^ melanoma cells (palbociclib treatment for 8 days) by induction of senolytic cell death, but not palbociclib-induced cell cycle arrested cells (palbociclib treatment for 1 day) (Fig 2B). While palbociclib or CB-839 alone induced marginal cell death, the combination did not enhance the effects in Nras^Q61^ cell lines unlike in Braf^WT^Nras^WT^ and Braf^V600E^ melanoma cell lines (Fig 2B). The treatments had a minimal impact on primary human melanocytes, indicating a low level of toxicity associated with this therapeutic approach (Fig 2B). Morphological analysis showed the damaged cells only after combined treatment of CDK4/6i and GLS1i in Braf^V600E^ melanoma cells (Sup Fig 2A). Consistent results were obtained using another GLS1/KGA inhibitor BPTES (sup Fig 2B), adding rigor and reproducibility to our results. We also assessed the remaining senescent cells after CB-839 treatment by senescence-associated β galactosidase (SA-βgal) assay. Palbociclib induces senescence in approximately 80% of cells consistent with our previous study ^27^ while CB-839 treatment did not induce senescence (Fig 2C-D). Although SA-βgal positive cells remained on the plate after the combined treatment of palbociclib and CB-839, the SA-βgal positivity was significantly diminished (Fig 2C-D). To determine the long-term efficacy of the combination, we performed a clonogenic colony formation assay. We treated skmel28 cells with palbociclib, CB-839 or both and then observed colony outgrowth for 35 days after the treatments. Although palbociclib or CB-839 alone suppressed colony formation, certain cells that evaded the treatments started growing (Fig 2E). However, the combination of palbociclib and CB-839 exhibited long-term suppression of colony formation by inducing cell death in senescent cells, which could potentially mitigate tumor recurrence, a significant challenge in cancer treatment, by eliminating therapy-induced senescent dormant cells (Fig 2E). Consistently, other FDA-approved CDK4/6 inhibitors for breast cancer such as abemaciclib and ribociclib induced senescence and KGA upregulation (Sup Fig 3A-B). This upregulation renders Braf^V600E^ melanoma cells susceptible to CB-839 as we observed in cells using palbociclib and CB-839 (Sup Fig 3C). Induction of senolysis was independent of DNA damage response as the combination of palbociclib and CB-839 did not induce ɣ-H2AX, a marker for DNA damage (Sup Fig 4). Together these results indicate that CDK4/6i-induced senescent cells upregulate KGA and additional treatment of GLS1i induces senolysis in senescent melanoma cells, except in Nras mutant melanoma cells. These findings may partially explain the lack of efficacy observed in the outcomes of a phase 1b/2 clinical trial (clinicaltrials.gov- NCT03965845) evaluating CB-839 and palbociclib in patients with solid tumors harboring Ras mutation. It may be necessary to redirect the clinical application of GLS1i, which is currently utilized to target proliferating cancer cells in clinical trials, toward the targeting of senescent cells instead.

**Figure 2.**
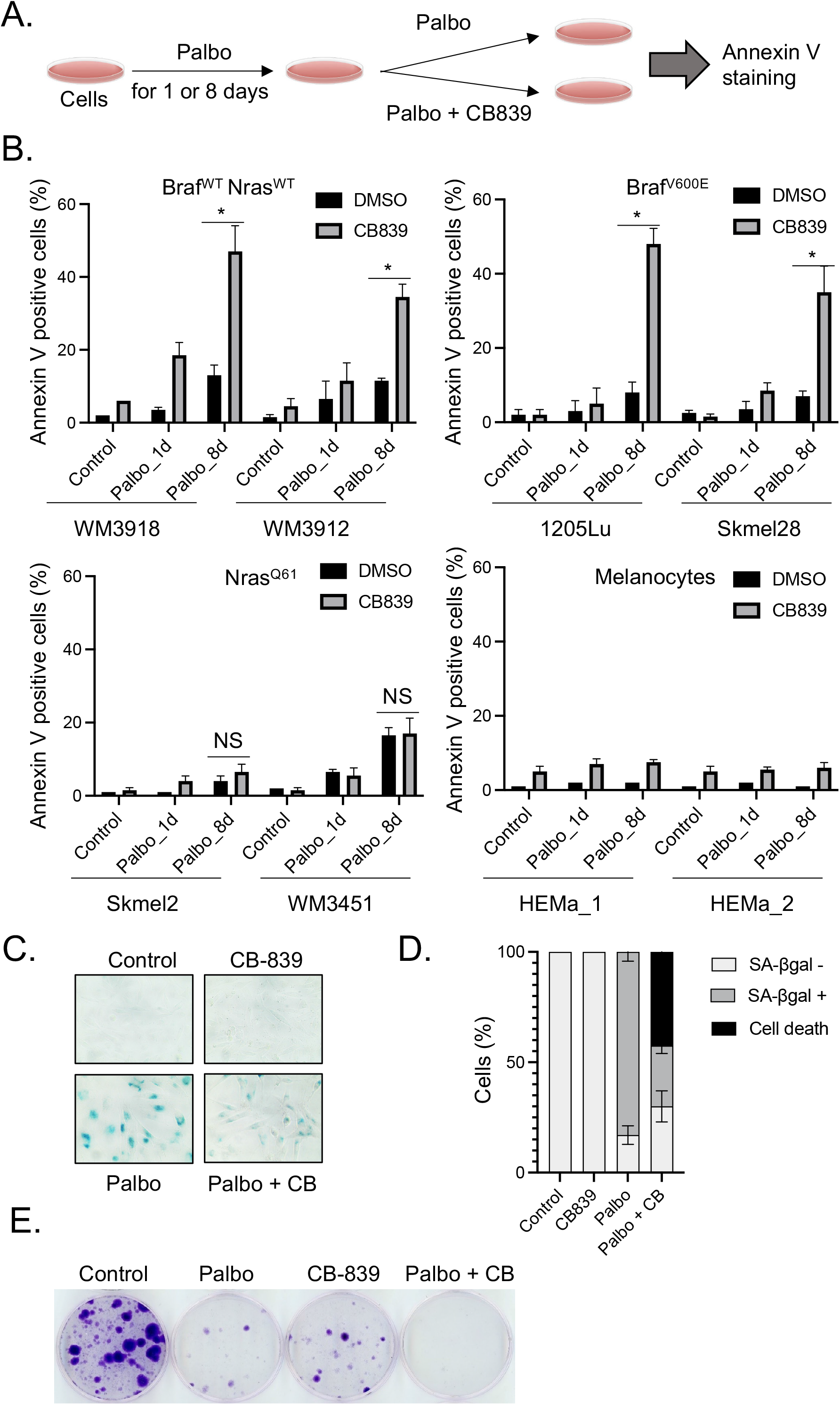
CB-839 induces senolysis in palbociclib-induced senescent cells. (A) Scheme for experimental schedule. (B) WM3918 (Braf^WT^Nras^WT^), WM3912 (Braf^WT^Nras^WT^), 1205Lu (Braf^V600E^Nras^WT^), Skmel28 (Braf^V600E^Nras^WT^), Skmel2 (Braf^WT^Nras^Q61R^), WM3451 (Braf^WT^Nras^Q61K^), HEMa_1 (normal human melanocytes) and HEMa_2 (normal human melanocytes) were treated with DMSO (Control) or palbociclib (1μM) for 1 day or 8 days followed by CB-839 (50μM) for an additional 3 days. Harvested cells were subjected to Annexin V staining and analyzed by flow cytometry. Data represent mean ± SD, * *p* < 0.05 (two-tailed Student’s t- test; n = 3), NS indicates non-significant (n = 3). (C) 1205Lu cells treated with DMSO (Control) or palbociclib (1μM) for 1 day or 8 days followed by CB-839 (50μM) for an additional 3 days were subjected to SA-βgal assay. (D) Quantification of SA-βgal positive cells from (C) was shown; data represent mean ± SD. (E) Skmel2 cells were treated with DMSO (Control), palbociclib (0.25 μM) for 8 days, CB-839 (50 μM) for 3 days, or palbociclib (0.25 μM) for 8 days followed by CB- 839 (50μM) for an additional 3 days. Colony outgrowth was stained with crystal violet after culturing cells for 35 days, including the treatment period.

### Knockdown of KGA with palbociclib induces senolysis in Braf^V600E^ melanoma cells

The above results demonstrated the potential utilization of the combination of CDK4/6i and GLS1i in targeting Braf^V600E^ melanoma. To assess the importance of upregulated GLS1 in CDK4/6i-induced Braf^V600E^ senescent cells and the specificity of the impact of CB-839, we conducted knockdown experiments. We introduced doxycycline-inducible shKGA, a main isoform of GLS1, in 1205Lu and Skmel28 Braf^V600E^ melanoma cells using two different shRNA hairpins (Fig 3A; the top band represent the KGA expression) and assessed cell death by Annexin V staining following treatment of them with palbociclib for 8 days and doxycycline for an additional 5 days. We demonstrated that GLS1 knockdown induced senolysis in palbociclib-induced senescent cells (Fig 3B), consistent with CB-839 treatment. To clarify the importance of KGA upregulation in palbociclib-induced senescent cells, we performed the same experiment using a doxycycline- inducible shGAC contract (Sup Fig 5A; the bottom band represents the GAC expression). Annexin V staining revealed no significant difference in cell death induction between treatments with palbociclib alone and the combination of palbociclib with doxycycline (Sup Fig 5B).

**Figure 3.**
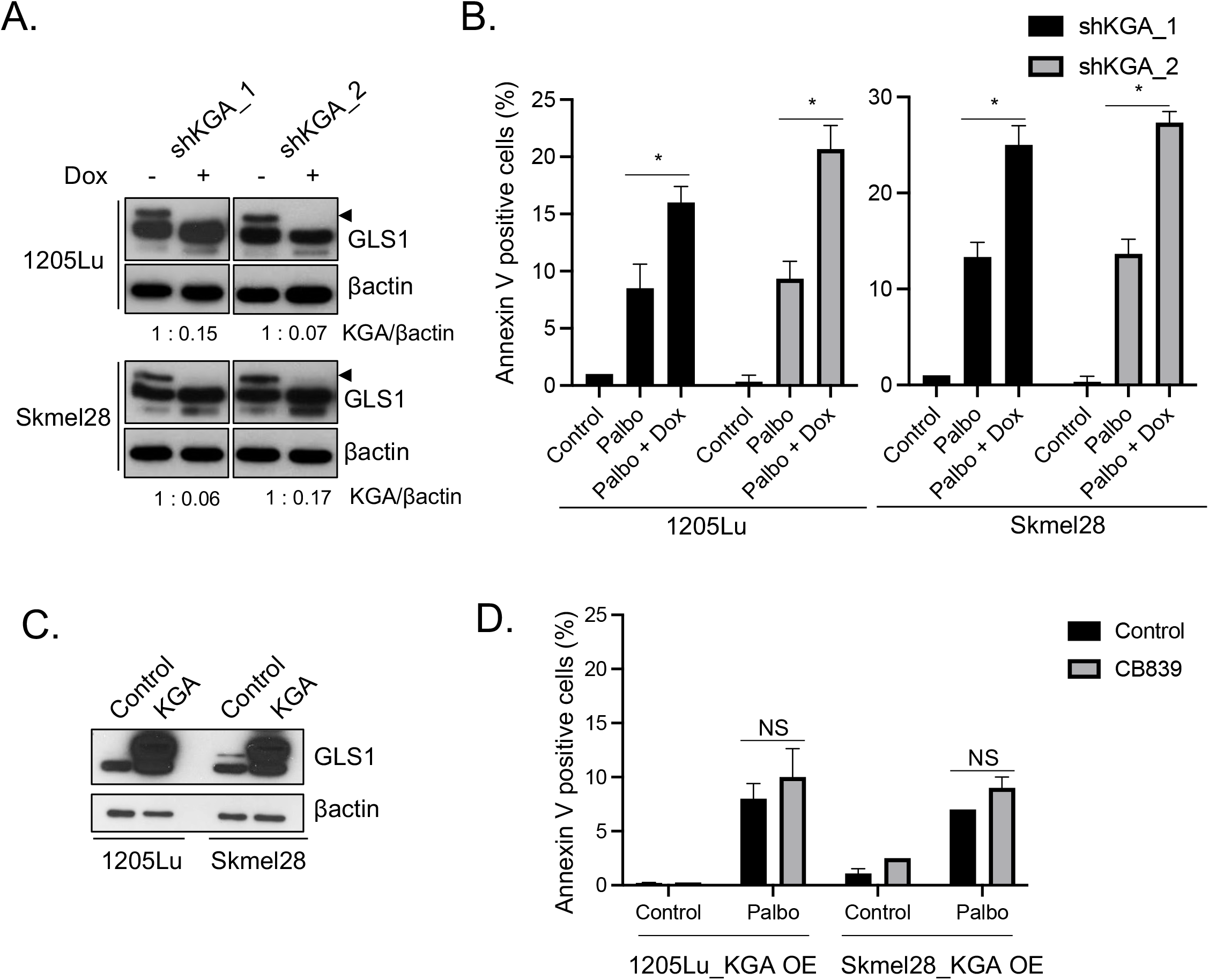
Knockdown of GLS1 induces senolysis in palbociclib-induced senescent cells. (A) 1205Lu and Skmel28 cell lines, transduced to stably express inducible short hairpin RNAs targeting KGA (shKGA_1 and shKGA_2), were treated with or without doxycycline (100 ng/ml) for 5 days and subjected to western blot for GLS1 and βactin. Arrowheads indicate KGA expression. (B) 1205Lu and Skmel28 cells from (A) were treated with DMSO (Control), palbociclib (1μM) for 13 days, or palbociclib (1μM) for 8 days followed by CB-839 for an additional 5 days. Harvested cells were subjected to Annexin V staining and analyzed by flow cytometry. Data represent mean ± SD, * *p* < 0.05 (two-tailed Student’s t-test; n = 3). (C) 1205Lu and Skmel28 cells introduced with overexpression of KGA were subjected to western blot for GLS1 and βactin. (D) 1205Lu and Skmel28 cells introduced with overexpression of KGA were treated with DMSO (Control) or palbociclib (1μM) for 8 days followed by CB-839 (50μM) for an additional 3 days. Harvested cells were subjected to Annexin V staining and analyzed by flow cytometry. Data represent mean ± SD, NS indicates non-significant (n = 3).

We next asked whether enforced expression of KGA enhances CB-839 mediated senolysis induction after treatment of palbociclib in Braf^V600E^ melanoma cells. We introduced KGA expression in 1205Lu and skmel28 Braf^V600E^ melanoma cells (Fig 3C) and then treated them with palbociclib for 8 days and CB-839 for an additional 3 days. Cell death was assessed by Annexin V staining. We demonstrated that although only palbociclib induced senolysis, additional CB-839 treatment did not enhance the efficacy of palbociclib in KGA overexpressing cells (Fig 3D). This indicates that the upregulated endogenous expression of KGA in palbociclib-induced senescent cells may suffice for cell survival, and further augmentation of KGA expression may not be essential. Together these results indicate that the upregulation of endogenous GLS1 (specifically KGA expression) by CDK4/6i is required for induction of cell death by GLS1i or GLS1 knockdown whereas enforced expression of KGA did not enhance its efficacy in Braf^V600E^ melanoma cells.

### CDK4/6i and GLS1i overcome vemurafenib resistance in Braf^V600E^ melanoma cells

To align with current clinical practice in melanoma management, we subsequently extended the Braf^V600E^ inhibitor (Brafi)-resistant population and assessed the efficacy of senolytic therapy using CDK4/6i and GLS1i in Brafi vemurafenib-resistant Braf^V600E^ melanoma cells. Brafi has been used as front-line therapy for Braf^V600E^ melanoma, but resistance to Braf^V600E^ inhibitor has been frequently observed clinically and is problematic for treating melanoma ^43^. We used two vemurafenib-resistant Braf^V600E^ melanoma cell lines derived from 1205Lu and 983B that we established previously ^27^. We first assessed the expression of KGA after treatment with palbociclib in vemurafenib-resistant (VR) Braf^V600E^ melanoma cells. We treated VR cells with palbociclib for 1 day or 8 days and harvested cells to assess KGA expression. We demonstrated that KGA is significantly induced after treatment of palbociclib for 8 days while 1-day treatment of palbociclib marginally induced KGA expression (Fig 4A). Upregulated KGA was further verifiedd by western blot (Fig 4B; the top band represents KGA expression). We next treated VR Braf^V600E^ melanoma cells with palbociclib for 1 day or 8 days, followed by CB-839 for an additional 3 days to assess apoptosis (as described in Fig 2A). We demonstrated that GLS1i induced senolysis in palbociclib-induced senescent cells, but not in cells without palbociclib treatment or palbociclib-induced cell cycle arrested cells (Fig 4C), consistent with results observed using vemurafenib sensitive Braf^V600E^ melanoma cells. Morphological analysis of 1205 VR cells showed that palbociclib induced senescence-like morphology, whereas the combination effectively damaged cells (Sup Fig 6). Together these findings suggest that the senolytic treatment combining CDK4/6i and GLS1i is effective for Braf^V600E^ melanoma cells, including those that have developed resistance to vemurafenib. This data demonstrates in vitro proof-of-concept that combination senolytic therapy may be a viable treatment option for patients with resistant melanoma and warrants further clinical trials to evaluate its efficacy.

**Figure 4.**
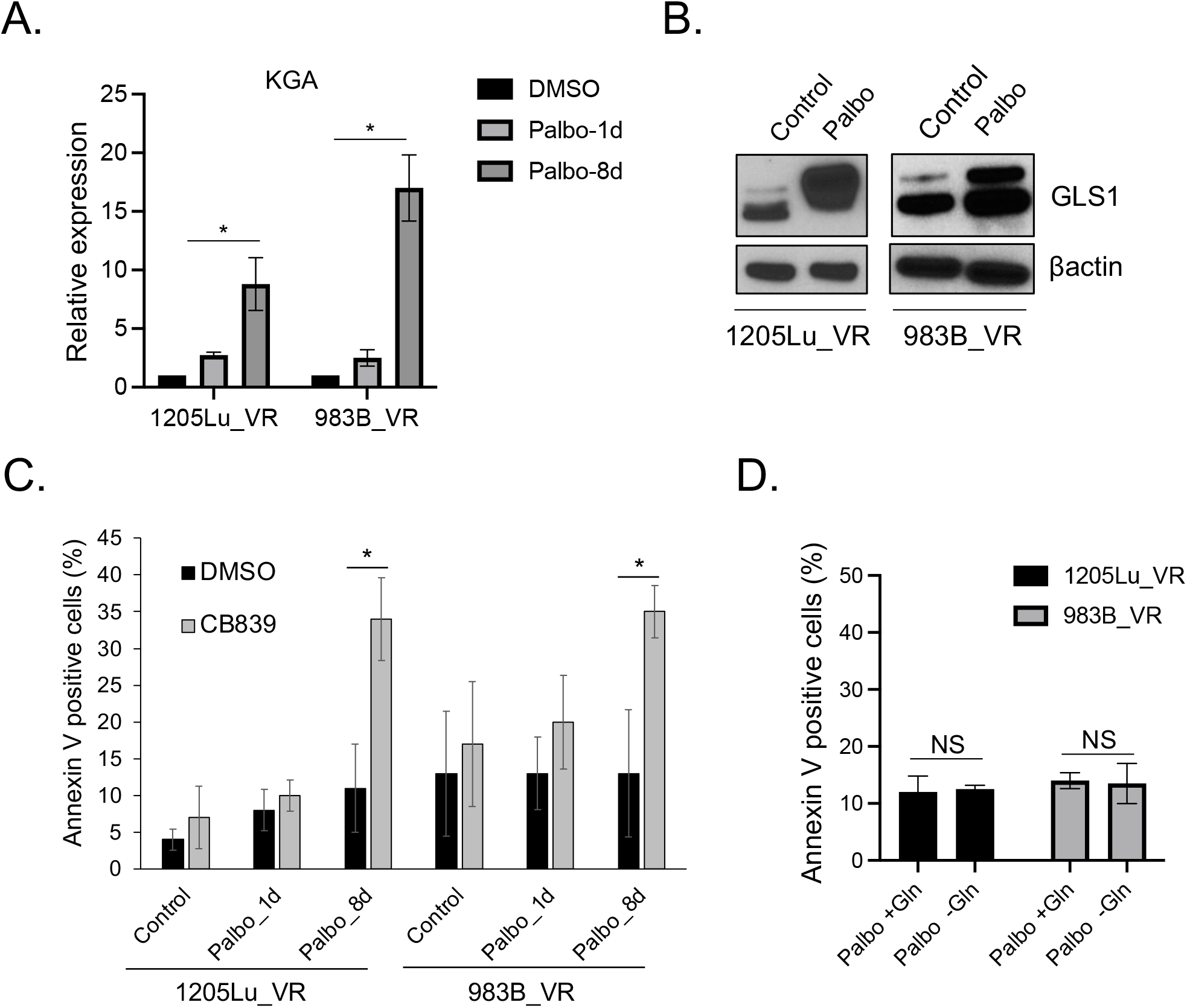
CDK4/6i and GLS1i induces senolysis in vemurafenib-resistant melanoma cells. (A) Vemurafenib-resistant cells derived from 1205Lu and 983B (1205Lu_VR and 983B_VR) treated with DMSO or palbociclib (1μM) for 1 day or 8 days were subjected to RT-qPCR analysis for KGA expression. Data were normalized by GAPDH and represent mean ± SD, * *p* < 0.05 (two- tailed Student’s t-test; n = 3). (B) 1205Lu_VR and 983_VR cells treated with DMSO (Control) or palbociclib (1μM) for 8 days were subjected to western blot for GLS1 and βactin. (C) 1205Lu_VR and 983B_VR cells were treated with DMSO (Control) or palbociclib (1μM) for 1 day or 8 days followed by CB-839 (50μM) for an additional 3 days. Harvested cells were subjected to Annexin V staining and analyzed by flow cytometry. Data represent mean ± SD, * *p* < 0.05 (two-tailed Student’s t-test; n = 3). (D) 1205Lu_VR and 983B_VR cells were treated with palbociclib (1μM) for 8 days and then cultured in a medium with or without glutamine for an additional 3 days. Harvested cells were subjected to Annexin V staining and analyzed by flow cytometry. Data represent mean ± SD, NS indicates non-significant (n = 3).

**Figure 5.**
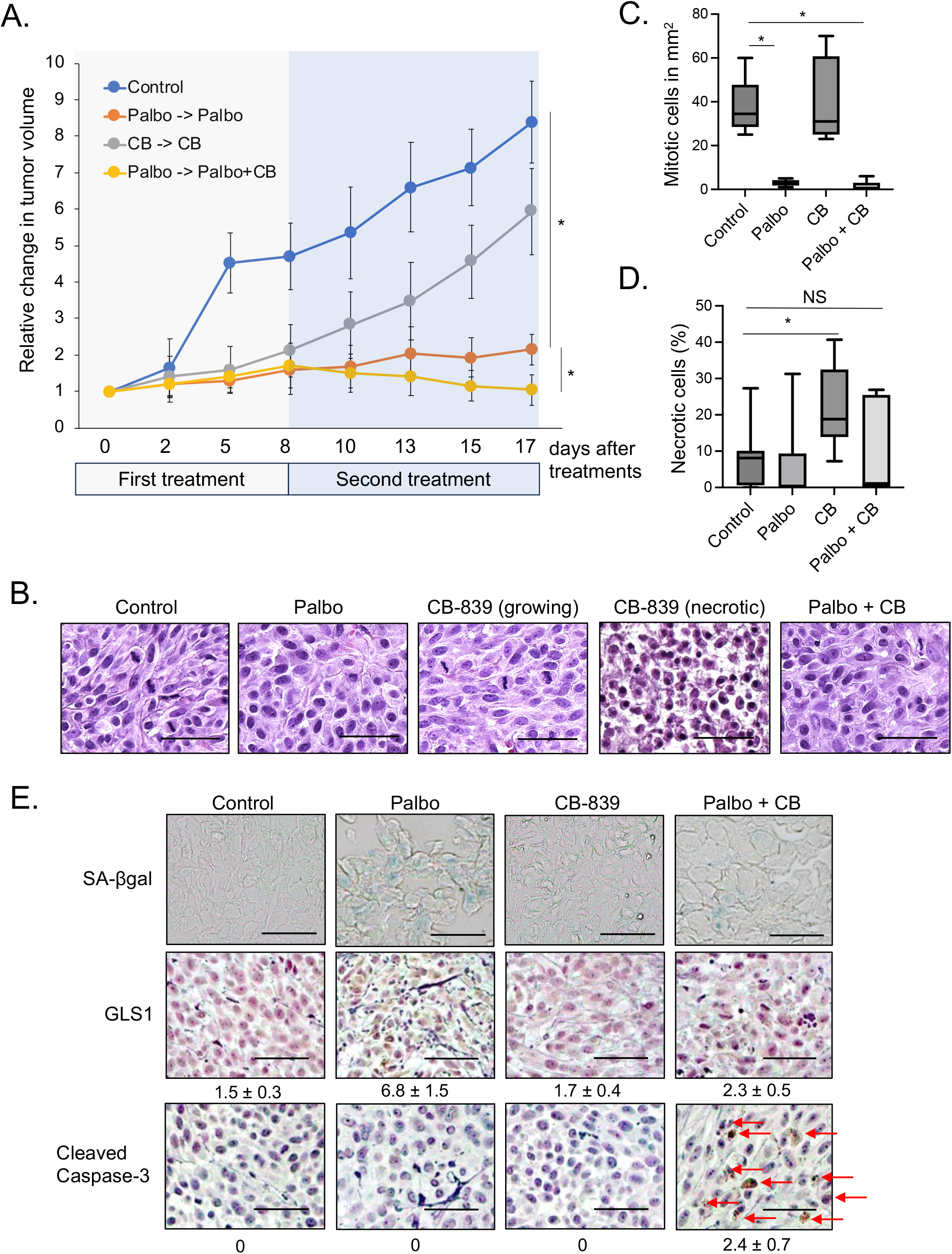
CDK4/6i and GLS1i induces senolysis *in vivo*. (A) A total of 1 × 10^6^ cells of 1205Lu_VR cells were subcutaneously injected into 6-week-old SCID mice. Tumor-inoculated mice were treated with palbociclib (90 mg/kg) by oral gavage or CB-839 (200 mg/kg) by oral gavage after tumors reached 5 mm in diameter. After 8 days, a combination of palbociclib (90 mg/kg) and CB-839 (200 mg/kg) was given by oral gavage for an additional 9 days. In each group, comprising 10 tumors, tumor volumes were measured every 2 days. Data represent means ± SD, * *p* < 0.05 (two-tailed Student’s t-test; n = 10). (B) Representative images of H&E staining for harvested tumors from (A). The scale bar indicates 50μm. (C) The number of mitotic cells was enumerated within a representative 1 mm^2^ area in each image. Data represent median and interquartile range (IQR), * *p* < 0.05 (two-tailed Student’s t- test; n = 10). (D) The proportion of necrotic cells was quantified in each H&E-stained image. Data represent median and interquartile range (IQR), * *p* < 0.05 (two-tailed Student’s t-test; n = 10). NS indicates non-significant (n = 10). (E) Representative images of SA-βgal staining and IHC staining for GLS1 and cleaved Caspase-3 in the tumors from (A). The numbers indicate H scores by IHC scoring (see Materials and Methods) from 10 tumors. The scale bar indicates 50μm. The red arrow indicates Cleaved Caspase-3 positive cells.

**Figure 6.**
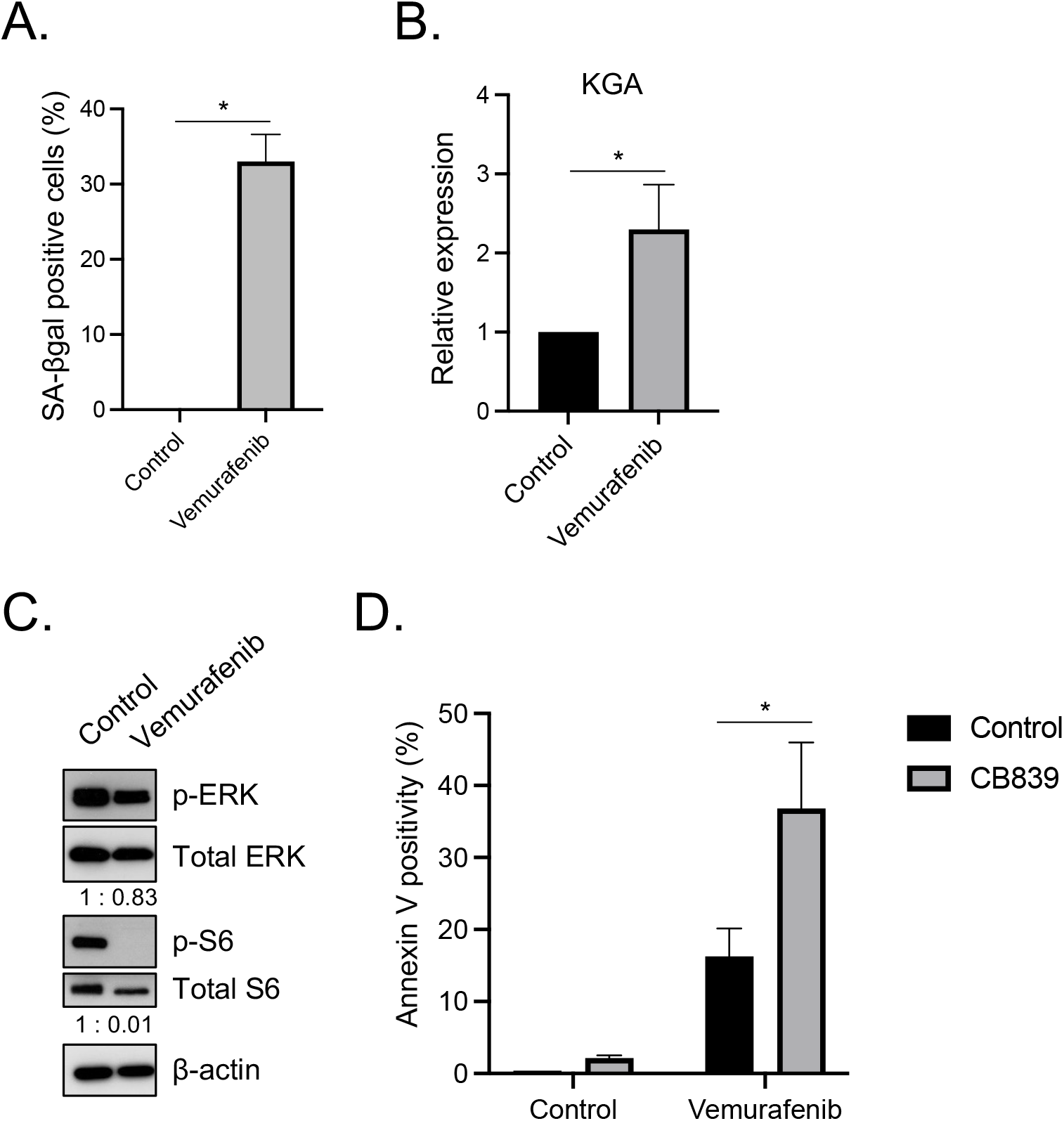
GLS1i induces senolysis in vemurafenib-induced senescent cells. (A-B) 1205Lu cells treated with DMSO (Control) or vemurafenib (0.5 μM) for 8 days were subjected to SA-βgal assay (A) and RT-qPCR analysis for KGA expression (B). RT-qPCR data (B) were normalized by GAPDH and represent mean ± SD, * *p* < 0.05 (two-tailed Student’s t-test; n = 3). (C) 1205Lu cells treated with DMSO (Control) or vemurafenib (0.5 μM) for 8 days were subjected to western blot for p-ERK, total ERK, p-S6, total S6 and βactin. (D) 1205Lu cells were treated with DMSO (Control) or vemurafenib (0.5 μM) for 8 days followed by CB-839 (50μM) for an additional 3 days. Harvested cells were subjected to Annexin V staining and analyzed by flow cytometer. Data represent mean ± SD, * *p* < 0.05 (two-tailed Student’s t-test; n = 3).

The above results have shown that induction of GLS1 expression (specifically KGA expression) by palbociclib in senescent cells is critical for successful targeting by GLS1i. Consistently, CDK4/6i increases mitochondrial metabolism in Braf^V600E^ melanoma cells ^36^ and pancreatic cancer ^35^. We asked whether palbociclib-induced senescence represents glutamine dependence, a state where cancer cells rely on glutamine for survival. We treated VR melanoma cells with palbociclib for 8 days and cultured them with medium lacking glutamine for an additional 3 days to assess cell death by Annexin V. We demonstrated that, unlike CB-839, glutamine depletion did not provoke senolysis in palbociclib-induced senescent cells (Fig 4D). This suggests that CDK4/6i-induced senescent cells rely on the upregulation of KGA expression rather than on glutamine itself for survival in Braf^V600E^ melanoma cells. Moreover, this indicates that KGA induction in senescent cells is critical target for GLS1i to provoke senolysis.

### GLS1 inhibitor induces senolysis in vemurafenib-resistant melanoma cells *in vivo*

To assess the efficacy of the combination of palbociclib and CB-839 *in vivo,* we performed a melanoma cell xenograft experiment. 1205Lu Braf^V600E^ VR melanoma cells were injected subcutaneously into 8-week-old SCID mice. We treated mice with either palbociclib alone for 17 days, CB-839 alone for 17 days, or pre-treatment of palbociclib for 8 days followed by the combination for an additional 9 days (total 17 days) after tumors had developed 5 mm in diameter (Fig 5A). While a single treatment of palbociclib suppressed melanoma cell proliferation through senescence induction as demonstrated previously ^27^, the combined treatment of palbociclib and CB-839 reduced tumor burden further compared to the palbociclib-alone treatment group (Fig 5A). Mouse body weights after treatment were unchanged (Sup Fig 7). Treatment of CB-839 alone is less effective in suppressing cell proliferation compared with palbociclib alone or a combination group (Fig. 5A). H&E staining revealed that the mitotic index, indicative of cell proliferation, was markedly reduced in palbociclib-alone and combination-treatment groups compared to the control group (Fig 5B-C). Despite the mitotic index of tumors treated with CB-839 being comparable to that of the control group, an increase in necrotic cell occurrence was observed exclusively in the CB-839-alone treated tumors (Fig 5B and D). The simultaneous presence of proliferating and necrotic cells provides insight into the modest inhibition of tumor growth by CB-839 treatment, even when the number of mitotic cells aligns with that of the control group. We further characterized the tumors following the various treatments. Using the SA-βgal assay we found that tumors treated with palbociclib-alone exhibited senescent cells and tumors treated with the combination of palbociclib and CB-839 showed less SA-βgal positivity than palbociclib-alone treated cells (Fig 5E top panel), suggesting CB-839 treatment effectively eliminated palbociclib-induced senescent cells *in vivo*. Further investigation using IHC staining revealed that GLS1 expression is upregulated in palbociclib-induced senescent cells, however, this upregulation was negated when tumors were concurrently treated with palbociclib and CB-839 (Fig 5E middle panel). Consistent with our *in vitro* results, we observed apoptosis by staining for cleaved Caspase- 3 expression in tumors treated with palbociclib and CB-839 (Fig 5E bottom panel). This staining indicated that although CB-839 itself has an impact on the induction of tissue necrosis, reflecting an attractive target for proliferating cancer cells, and currently being evaluated in many clinical trials for different types of cancers. However, senolytic therapy induced by the combination of palbociclib and CB-839 significantly reduces tumor burdens, which may be a more attractive strategy compared with targeting proliferating cancer cells by CB-839. Taken together these *in vivo* results strongly support our concept that induction of senolysis by targeting GLS1 in palbociclib-induced senescent cells is highly effective. Furthermore, these data indicate that senolytic therapy may be a potential therapeutic solution to address the clinical challenge of Braf^V600E^ inhibitor resistance in treating Braf^V600E^ melanoma. Further optimization of drug dosages and treatment schedules will be needed to enhance its potential efficacy.

### GLS1 inhibitor induces senolysis in vemurafenib-induced senescent cells

The results thus far demonstrated that targeting upregulated GLS1 by a GLS1 inhibitor in CDK4/6i-induced senescent cells is an attractive strategy for treating Braf^V600E^ melanoma. While we demonstrated that CDK4/6i induces senescence in Braf^V600E^ melanoma, the previous study has shown that a low concentration of vemurafenib induces senescent features in Braf^V600E^ melanoma cells ^44^. We next tested whether senescent cells induced by other therapeutics like vemurafenib are susceptible to GLS1i. We treated Braf^V600E^ 1205Lu melanoma cells with vemurafenib for 8 days and assessed senescence induction by SA-βgal assay and the expression of KGA by QPCR. Vemurafenib treatment for 8 days induced senescence (Fig 6A) and KGA expression (Fig 6B). The inhibition of the MAPK pathway was confirmed by the reduced phosphorylation of ERK (Fig 6B). Additionally, the phosphorylation of S6, a target of ERK, was also diminished (Fig 6C). This suggests that a low dose of vemurafenib suppresses downstream signaling. We next administered vemurafenib to 1205Lu melanoma cells for 8 days and then treated them with CB-839 for an additional 3 days. Cell death was then assessed using Annexin V, as detailed in Figure 2A. We demonstrated that this sequential treatment effectively induced the death of vemurafenib-triggered senescent cells (Fig 6D). Together, these data suggest that GLS1 represents a strategic molecular target in senescent cells with elevated KGA, induced not only by CDK4/6i but also by other cytostatic drugs such as a low concentration of vemurafenib. The potential to optimize senolytic therapeutic strategies through the use of GLS1i is significant, contingent upon the identification of pharmacological combinations that elicit a more robust and enduring senescent state, concomitant with KGA expression.

## Discussion

Therapeutic inhibition of CDK4/6 has been recognized as a potent anticancer strategy. Selective CDK4/6 inhibitors (CDK4/6i), such as palbociclib, abemaciclib and ribociclib have received FDA approval for the treatment of estrogen receptor-positive (ER+)/HER2-negative breast cancer. While CDK4/6 inhibition has been shown to effectively suppress tumor proliferation by inducing senescence in both *in vitro* and *in vivo* preclinical melanoma models, its efficacy has not been replicated in clinical trials for melanoma. Although the development of acquired resistance to CDK4/6i presents a challenge, recent evidence suggests that senescence can paradoxically promote tumorigenesis and recurrence by secreting various inflammatory cytokines ^5–8^. This underscores the necessity to innovate and develop novel senolytic agents capable of eradicating senescent cells induced by current therapeutic interventions, thereby circumventing the onset of resistance and mitigating the risk of neoplastic recurrence.

In this study, we assessed the expression of Glutaminase 1 (GLS1) in CDK4/6i-induced senescent melanoma cells and tested the applicability of targeting GLS1 to eliminate those senescent cells by GLS1 inhibition-mediated acidosis induction, given a recent study describing GLS1 as an essential gene supporting senescent cells for survival by neutralizing acidic conditions through ammonia production induced by upregulated GLS1 expression ^37^. We demonstrated that prolonged treatment of CDK4/6i upregulates GLS1 expression in Braf^V600E^ melanoma cells, which is consistent with recent studies demonstrating that CDK4/6i reprograms mitochondria metabolism ^35, 36^. This mitochondrial reprogramming may occur through GLS1 regulation since the expression of glutamine transporters such as SLC1A5 (ASCT2) is unchanged after treatment of palbociclib (Sup Fig 8). We further demonstrated that targeting GLS1 using a specific glutaminase inhibitor (CB-839) eliminates Braf^V600E^ senescent cells induced by either CDK4/6i or vemurafenib.

Importantly, the synergistic pharmacological intervention comprising palbociclib and CB-839 retains its efficacy against Braf^V600E^ melanoma cells exhibiting acquired resistance to vemurafenib, as evidenced by both *in vitro* and *in vivo* studies. Although we demonstrated that tumor growth suppression by senolysis induction using a combination of CDK4/6i and GLS1i is durable using long-term colony formation assay (Fig 2E), subsequent research evaluating the lifespan of mice bearing Braf^V600E^ melanoma following treatment with CDK4/6i and GLS1i could yield a more accurate assessment of this therapeutic regimen. This approach provides a promising therapeutic alternative for the management of Braf^V600E^ and Braf^WT^Nras^WT^ and melanoma treatment, particularly for patients who have developed resistance to MAPK-directed therapies and consequently face a paucity of viable treatment options.

Statistically, Braf and Nras mutations are almost mutually exclusive according to cBioPortal. Our research has established that treatment with CDK4/6i does not upregulate KGA expression in melanoma cells harboring Nras mutations. Combined treatment with CB-839 in these cells did not exhibit synergistic effects. These observations may partially elucidate the clinical ineffectiveness of the combined CDK4/6i and GLS1i in treating solid tumors with Ras mutations (clinicaltrials.gov- NCT03965845). These are relevant to developing a clinical trial as we need to test Braf mutations only. Interestingly, recent work revealed that the combination of CDK4/6i and MEKi induces senescence in pancreatic ductal adenocarcinoma ^45, 46^ and suppressed tumor growth in Braf/MEK inhibitor-resistant melanoma ^47^. Furthermore, the triple combination of Braf, MEK and CDK4/6 has shown promising results ^48, 49^ and has been evaluated in a clinical trial (clinicaltrials.gov- NCT04720768). We also previously demonstrated that an mTORC1 inhibitor (everolimus) enhanced colony outgrowth after co-culture with palbociclib compared to palbociclib alone ^26^. Recent cumulative research suggests that further investigation to induce a more durable senescent state through such cytostatic drug combinations could advance senolytic therapy targeting upregulated GLS1. Moreover, the co-administration of GLS1i to existing treatment regimens may permit dosage reduction, potentially diminishing adverse side effects and enhancing patient tolerance in clinical trials.

CB-839 demonstrated limited efficacy as an antineoplastic drug, which was notably evident in human liver cancer cell lines with significant glutamine dependence. Consequently, its combined use with other pharmacological agents has been explored across various cancer types. For instance, 5-fluorouracil (5-FU) has been noted to augment the antitumor activity of CB-839 in PIK3CA mutant colorectal cancers ^50^, and the proteasome inhibitor carfilzomib has shown synergistic effects when used with CB-839 in multiple myeloma ^51^. Given that a range of treatments, including chemotherapy and radiotherapy, are known to induce cellular senescence, employing CB-839 could strategically target cells rendered senescent following these therapies. Indeed, senescence induction has been documented in response to 5-FU ^52^.

Collectively, our study elucidates the scientific underpinnings for employing GLS1i in oncology and opens avenues for repurposing CB-839, which has hitherto shown limited success in clinical trials. Strategizing to identify drug combinations that elicit senescence accompanied by KGA upregulation – targets amenable to GLS1i – represents an innovative therapeutic approach for cancer management. This strategy diverges from traditional methods that primarily aim to exploit the glutamine dependency of cancer cells to directly induce apoptosis. Further research to advance this novel senolytic strategy using CDK4/6i and GLS1i into clinical trials is anticipated, with a focus on targeting Braf^V600E^ melanoma, inclusive of variants with acquired resistance to Braf and/or MEK inhibitors.

## Acknowledgments

We would like to express our sincere gratitude to Dr. Thomas McCormick for his invaluable suggestions and insightful critiques during the preparation of this manuscript. We also would like to thank Ms. Mamuni Swain at the Athymic Animal & Xenograft Core for her excellent assistance of *in vivo* experiments. Our appreciation also extends to the staff and facilities of the Animal Resource Center, the CDDRCC Histology Core, and the Cytometry & Imaging Microscopy Core at Case Western Reserve University, as well as the Case Comprehensive Cancer Center. Their support and expertise were instrumental in the completion of our research.

## Author contributions

J.K., B.B., A.K., and A.Y. performed the experiments. K.H. analyzed the mitotic index and the proportion of necrotic cells within HE-stained sections derived from murine xenograft tumors.

A.M. provided insightful feedback on the projects. J.K. and A.Y. wrote the manuscript.

