## Supplemental figures for "Inhibition of glutaminase elicits senolysis in therapy-induced senescent melanoma cells"

### Supplemental Fig 1.

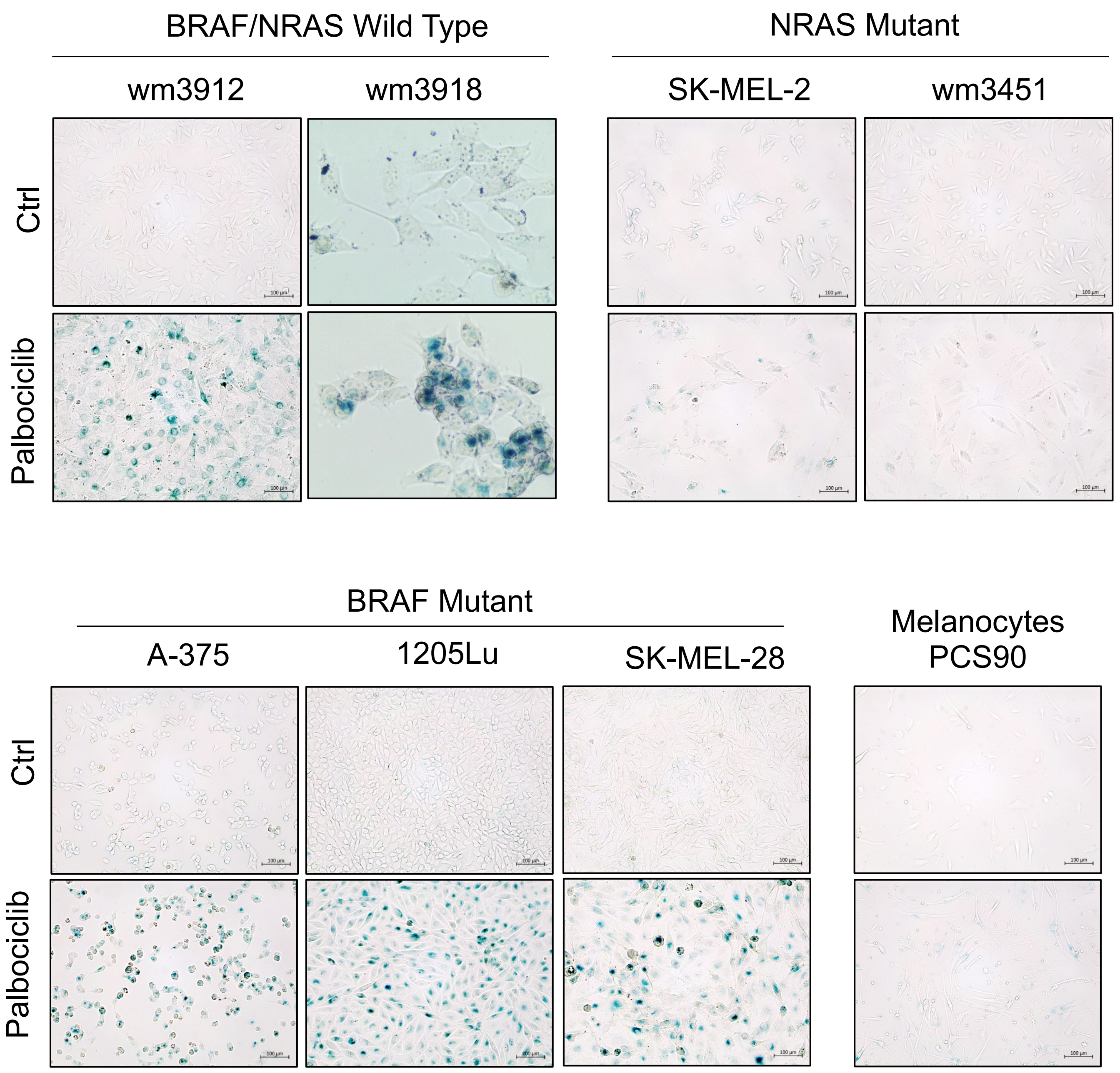


**Fig S1. Palbociclib induces senescence in Nras WT melanoa.**

WM3912 ($Braf^{WT}Nras^{WT}$), WM3918 ($Braf^{WT}Nras^{WT}$), A-375 ($Braf^{V600E}Nras^{WT}$), 1205Lu ($Braf^{V600E}Nras^{WT}$), Skmel28 ($Braf^{V600E}Nras^{WT}$), Skmel2 ($Braf^{WT}Nras^{Q61R}$), WM3451 ($Braf^{WT}Nras^{Q61K}$), and PCS90 (melanocytes) were treated with DMSO (Control) or palbociclib (1µM) for 8 days and subjected to SA-βgal staining. Representative images of SA-βgal staining are shown.

### Supplemental Fig 2.

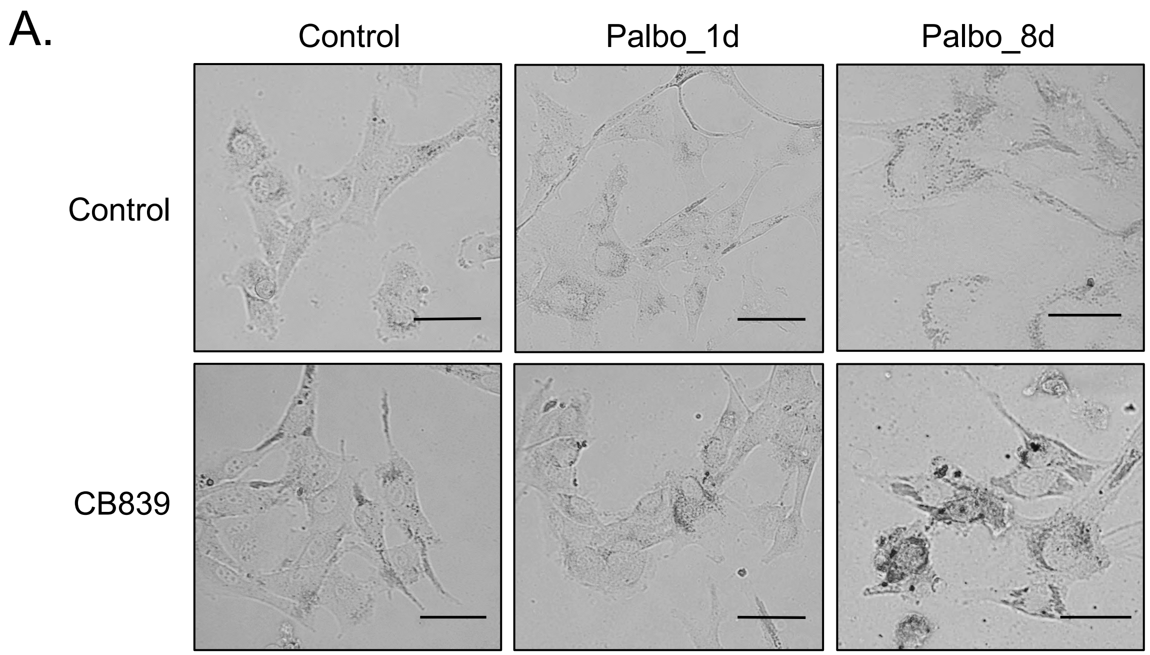


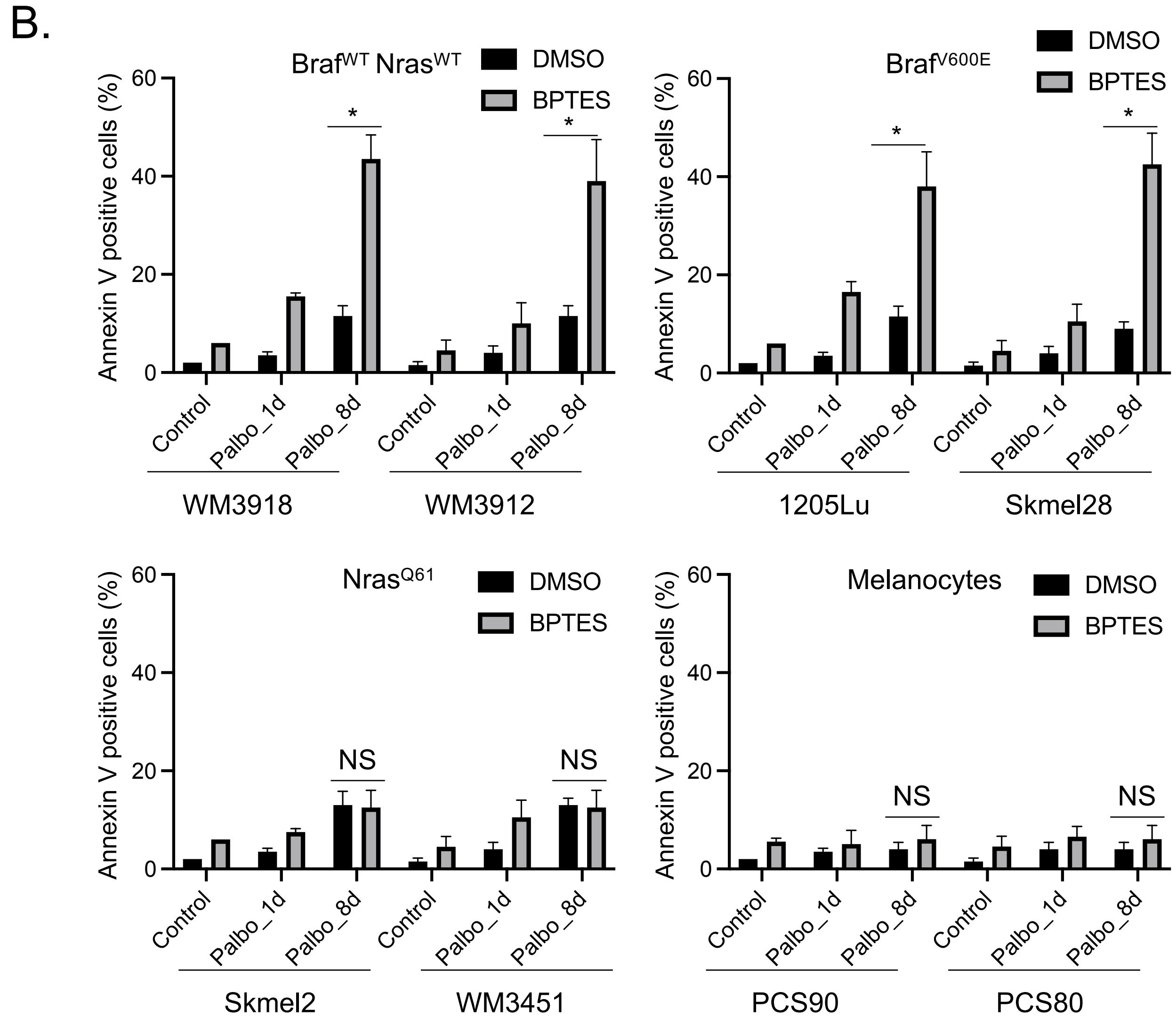

**Fig S2. GLS1i induces senolysis in palbociclib-induced senescent cells.**

(A) Skmel28 cells were treated with DMSO (Control) or palbociclib (1μM) for 1 day or 8 days followed by CB-839 (50μM) for an additional 3 days. Representative phase contrast images of each treatment were shown. Scale bar indicates 50μm. (B) WM3918 ($Braf^{WT}Nras^{WT}$), WM3912 ($Braf^{WT}Nras^{WT}$), 1205Lu ($Braf^{V600E}Nras^{WT}$), Skmel28 ($Braf^{V600E}Nras^{WT}$), Skmel2 ($Braf^{WT}Nras^{Q61R}$), WM3451 ($Braf^{WT}Nras^{Q61K}$), PCS90 (melanocytes) and PCS80 (melanocytes) were treated with DMSO (Control) or palbociclib (1μM) for 1 day or 8 days followed by BPTES (5μM) for an additional 3 days. Harvested cells were subjected to Annexin V staining and analyzed by flow cytometry. Data represent mean ± SD, * $p < 0.01$, NS non-significant.

### Supplemental Fig 3.

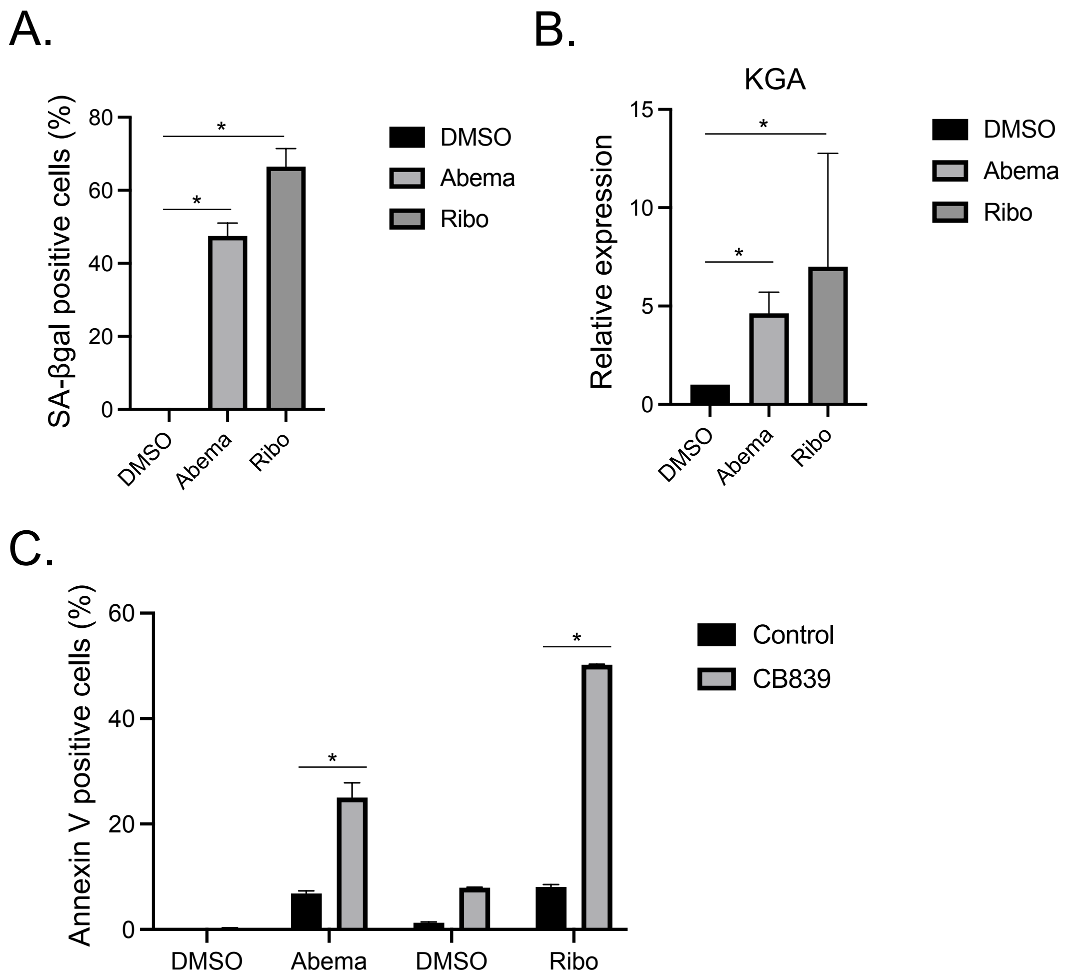


**Fig S3. GLS1i induces senolysis in CDK4/6i-induced senescent cells.** (A-B) 1205Lu cells were treated with DMSO (Control), abemaciclib (Abema) (1μM) or ribociclib (Ribo) (1μM) for 8 days. Harvested cells were subjected to SA-βgal assay (A), RT-qPCR analysis for KGA (B). Data were normalized by GAPDH and represent mean ± SD, * $p < 0.05$ (two-tailed Student's t-test; n = 3). (C) 1205Lu cells were treated with DMSO (Control), abemaciclib (1μM) or ribociclib (1μM) for 8 days followed by CB-839 (50μM) for an additional 3 days. Harvested cells were subjected to Annexin V staining and analyzed by flow cytometry. Data represent mean ± SD, * $p < 0.01$, NS non-significant.

### Supplemental Fig 4.

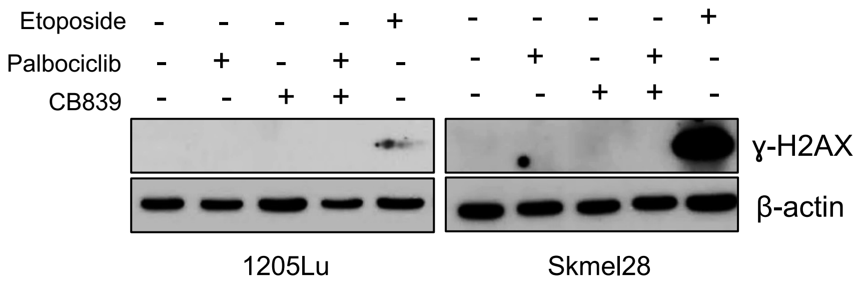


**Fig S4. Palbociclib and CB-839 do not induce DNA damage.**

(A) 1205Lu and Skmel28 cells were treated with DMSO (Control), palbociclib for 11 days, CB-839 for 3 days, or palbociclib (1μM) for 8 days followed by CB-839 (50 μM) for an additional 3 days. Etoposide treatment (10 μM) was used as a positive control. Harvested cells were subjected to western blot for ɣ-H2AX and β-actin.

### Supplemental Fig 5.

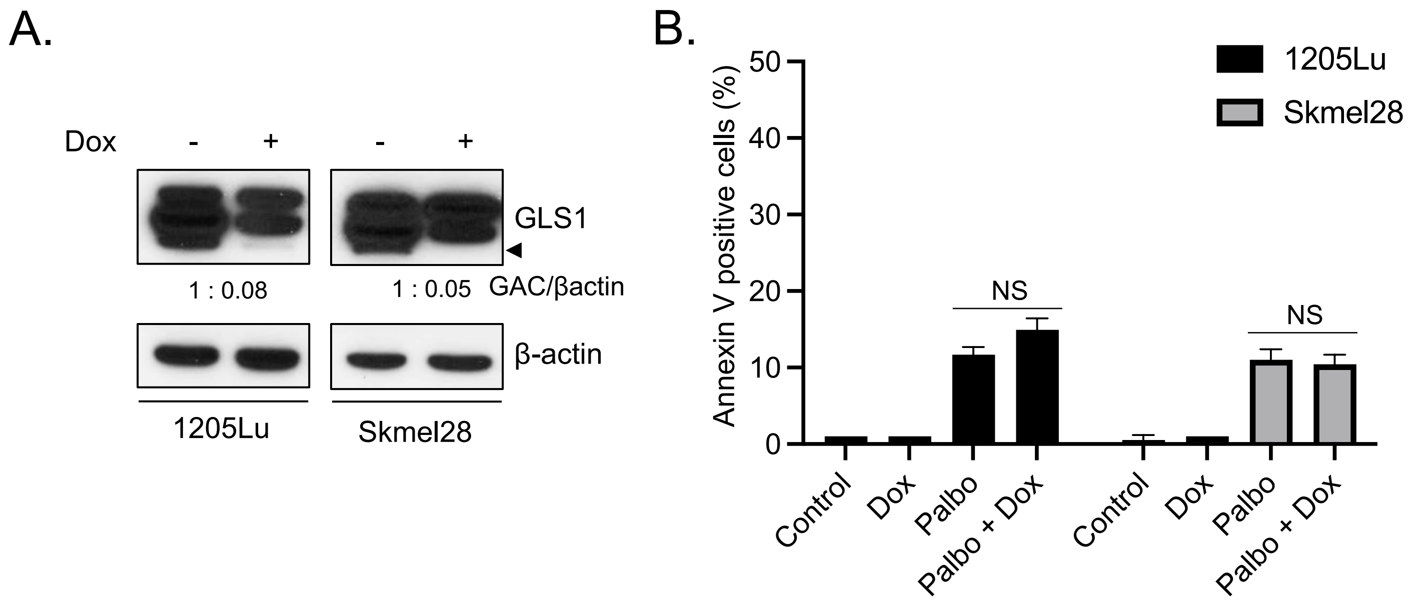


**Fig S5. ShGAC does not induce senolysis in palbociclib-induced senescent cells.**

(A) 1205Lu and Skmel28 cells were introduced with doxycycline-inducible shGAC. After selection, cells were treated with doxycycline (100 ng/ml) for 5 days. Harvested cells were subjected to western blot for GLS1 and β-actin. The bottom band indicates GAC. Arrowheads indicate GAC expression. (B) Cells from (A) were treated with palbociclib (1μM) for 13 days, doxycycline (100 ng/ml) for 5 days, or palbociclib (1μM) for 8 days followed by doxycycline (100 ng/ml) for an additional 5 days. Harvested cells were subjected to Annexin V staining and analyzed by flow cytometry. Data represent mean ± SD, NS non-significant.

### Supplemental Fig 6.

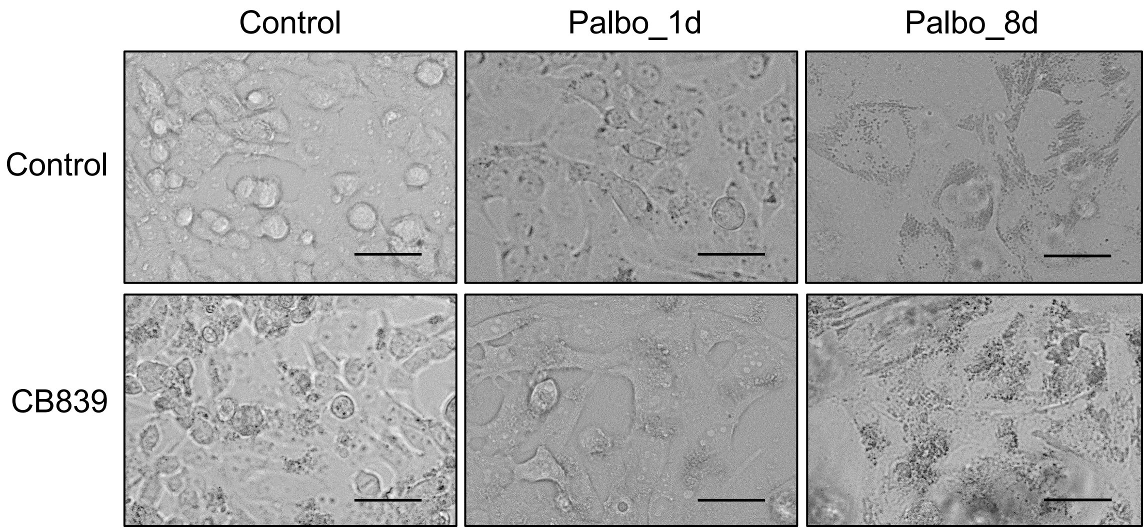


**Fig S6. GLS1i induces senolysis in palbociclib-induced vemurafenib-resistant senescent cells.**

1205Lu vemurafenib-resistant (VR) cells were treated with DMSO (Control) or palbociclib (1μM) for 1 day or 8 days followed by CB-839 (50μM) for an additional 3 days. Representative phase contrast images of each treatment are shown. Scale bar indicates 50μm.

### Supplemental Fig 7.

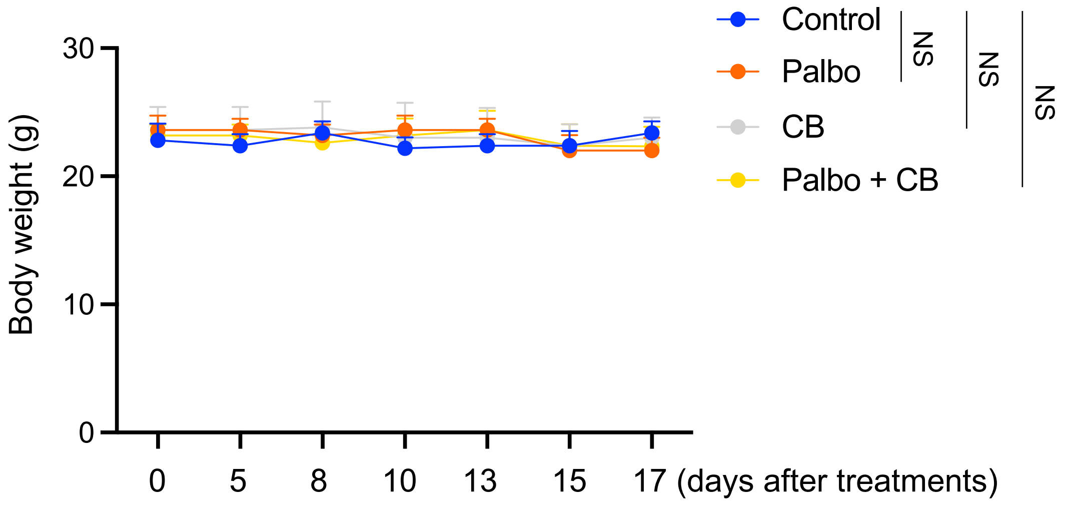


**Fig S7. mouse body weights are unchanged after treatments.**

Mouse body weights were measured at indicated time points in each mouse after treatments. NS indicates non-significant (n = 5).

### Supplemental Fig 8.

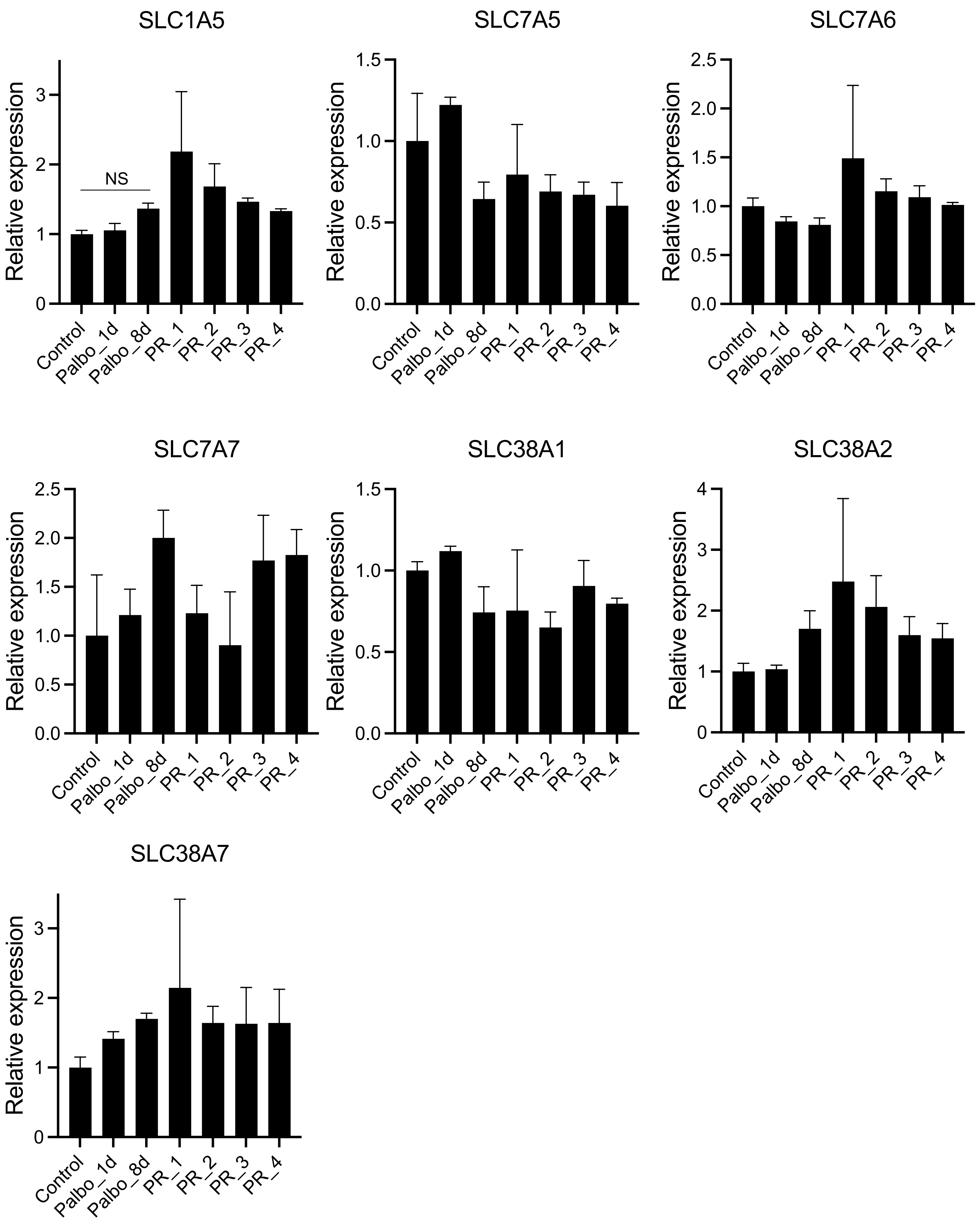

**Fig S8. Expression of glutamine transport is unchanged.**

(A) 1205Lu cells treated with DMSO (Control) or palbociclib (1μM) for 1 day or 8 days and 4 clones of 1205Lu palbociclib resistant cells (PR_1, PR_2, PR_3 and PR_4) were harvested and subjected to RNA-seq in a previous study (Yoshida et al, *Sci Adv* 2019). Data mining of RNA seq data for the expression of glutamine transporters is shown. Data represent mean ± SD.
